# TRACEDD: A Tool-grounded Reasoning and Agentic Coordination for Explainable Drug Design

**DOI:** 10.64898/2026.09.12.751167

**Authors:** Sarveswara Rao Vangala, Vishnu Vardhan Kasturi, Navneet Bung, Arijit Roy

## Abstract

Drug discovery depends on coordinated decisions across target validation, structure analysis, molecular design, developability assessment and synthetic feasibility, but current computational methods often operate as disconnected tools. Here, we introduce TRACEDD (Tool-grounded Reasoning and Agentic Coordination for Explainable Drug Design), a framework that makes three primary contributions: (1) It establishes a ‘tool-first’ multi agentic architecture where LLMs orchestrate validated computational tools rather than replace them, ensuring scientific rigor. (2) It implements a multi-agent system that mirrors expert discovery teams, enabling transparent and traceable decision-making through a Reason-Act-Observe loop. (3) It demonstrates an end-to-end workflow, from target validation to synthesis planning, that adaptively handles real-world data variability, such as the absence of experimental structures. The framework decomposes discovery into specialized agents for target validation, druggability assessment, molecular generation, lead optimization, ADMET evaluation, literature evidence integration and retrosynthesis, all operating through a Reason–Act–Observe workflow. Using JAK2 as a representative case, we show that the system can retrieve experimental protein structures, invoke AlphaFold when structures are unavailable, identify druggable pockets and perform de novo molecular generation. Known JAK2 inhibitors are used to define design hypotheses and guide reinforcement learning-based molecular generation, with docking scores/predicted pIC50 and other physicochemical/ADMET properties serving as reward and prioritization signals. The framework demonstrates a tool-first, reasoning-driven approach in which each major decision is linked to explicit tool invocation, intermediate evidence. By combining agentic orchestration with domain-specific computational tools, the system supports transparent, adaptable and human-verifiable molecular design workflows, providing a foundation for more reliable AI-assisted drug discovery.

## Introduction

Drug discovery is an iterative process of hypothesis generation, evidence integration and decision-making across structural biology, medicinal chemistry, pharmacokinetics, and synthetic feasibility. While recent advances in machine learning and artificial intelligence, together with physics-based methods, have substantially improved individual components of this pipeline including protein structure prediction,^1,2^ molecular generation,^3–10^ binding affinity estimation,^11,12^ ADMET prediction,^13,14^ and retrosynthesis planning^15^ these advances have largely emerged as standalone tools that operate in isolation. As a result, critical challenges remain in coordinating decisions across stages, propagating uncertainty and evidence, and integrating heterogeneous outputs into coherent, scientifically grounded design strategies.

Large language models (LLMs) have recently been proposed as general-purpose interfaces for scientific workflows, with early agentic frameworks demonstrating that LLMs can act as reasoning and control layers rather than as passive generators. In the Reason–Act–Observe (ReAct) paradigm, reasoning is tightly coupled to tool use, enabling the system to iteratively evaluate evidence, refine hypotheses, and guide subsequent decisions.^16^ By coupling reasoning with explicit actions and grounded observations, this approach reduces reliance on latent inference alone and improves both interpretability and robustness of AI-assisted decision-making.

Emerging evidence from scientific agent systems further suggests that LLMs are most effective when positioned as orchestration layers that coordinate established, domain-validated tools rather than attempting to replace them with end-to-end black-box predictors.^16^ Tool-aware, multi-agent frameworks have shown promise in biomedical applications by enabling conversational planning and cross-stage reasoning while preserving the rigor, trust, and familiarity of existing computational methods.^17,18^ This design choice aligns closely with how experts operate in practice, where confidence in results is derived from traceable evidence, validated models, and explicit reasoning rather than from predictive scores alone.

In parallel, the rapid expansion of high-quality protein structural data driven by experimental repositories and large-scale structure prediction methods have enabled structure-guided drug design even for targets lacking experimentally resolved structures.^1,2^ At the same time, generative chemistry platforms based on reinforcement learning now allow multi-objective optimization over potency, physicochemical properties, and synthetic accessibility.^3,7,9^ Yet these methods do not independently determine which structure should be used, whether a binding pocket is sufficiently druggable, which properties should govern generation, how conflicting property predictions should be reconciled or whether a proposed molecule has a plausible synthetic route. These are connected scientific decisions rather than isolated prediction tasks.

Here we present TRACEDD, a tool-grounded, multi-agent framework that represents early-stage drug design as a sequence of evidence-linked scientific decisions. A supervisory orchestrator decomposes a research objective into tasks addressed by role-specialized agents for target validation, druggability assessment, molecular generation, lead optimization, ADMET and bioactivity evaluation, literature evidence integration and retrosynthesis planning. The agents operate through a shared Reason–Act–Observe model in which scientific questions are translated into explicit tool calls, outputs are returned as structured observations and the accumulated evidence informs subsequent actions. TRACEDD follows a tool-first design principle. LLMs are used to plan tasks, select tools, integrate heterogeneous outputs and articulate the rationale connecting one decision to the next. Domain-specific computational methods remain the source of structural, chemical, pharmacological and synthetic predictions. The framework keep records of tool calling, intermediate outputs and decision rationales, thereby creating an auditable link between the initial scientific objective and the resulting design recommendations. This separation between orchestration and prediction is intended to limit unsupported inference while preserving the flexibility required to adapt a workflow as new evidence or constraints emerge.

We demonstrate the framework through connected use cases spanning target-structure identification/generation, evidence-linked hypothesis generation, programmatic preparation of protein structures, molecular generation against the target protein, property-directed lead optimization, candidate prioritization and retrosynthetic analysis. Rather than evaluating the platform solely by the number of molecules generated, we examine whether it can translate a high-level scientific objective into a traceable sequence of computationally grounded decisions. TRACEDD thus provides a framework for investigating how agentic AI can support reasoning-centered drug design while retaining human oversight, explicit provenance and compatibility with established scientific tools.

## Method

### Overall System Architecture

Early-stage drug discovery is formulated as a multi-agent system in which distinct stages of the workflow are decomposed into role-specialized computational agents operating over a shared execution state. These tasks span the core stages of medicinal chemistry and translational decision-making, including understanding and validating the biological target, assessing druggability and identifying viable binding opportunities, designing and optimizing molecular structures, evaluating developability and risk, validating novelty and prior art from existing literature, and assessing synthetic feasibility.

Each agent operates autonomously within its domain while remaining tightly coupled to the broader workflow. An agent reasons over its local scientific context, explicitly invokes external computational or data-driven tools, and produces structured, inspectable outputs accompanied by explicit reasoning, allowing subsequent steps to reuse, refine, or challenge upstream results. A supervisory orchestrator translates high-level objectives into coordinated workflows composed of sequential and parallel tasks. This agentic formulation closely mirrors how human discovery teams operate in practice where structural biologists, medicinal chemists, ADMET scientists, and synthetic chemists contribute specialized expertise while enabling tighter coordination, faster iteration, and persistent memory across design cycles. The intermediate evidence determines subsequent actions, enabling iterative refinement of hypotheses and design strategies. This approach preserves continuity across stages while maintaining clear traceability of decisions and their underlying evidence.

#### 2. Design Principles: ReAct + Tool-First Orchestration

The framework is structured around two complementary design principles: a Reason–Act–Observe (ReAct)^16^ execution paradigm and a tool-first orchestration strategy, which together enable grounded, interpretable, and iterative scientific decision-making.

Under the ReAct paradigm, each agent operates through an explicit reasoning loop in which intermediate hypotheses are formulated, translated into concrete actions via tool invocation, and subsequently updated based on observed outputs. This tightly coupled reasoning–action–feedback cycle ensures that decisions are continuously informed by empirical or computational evidence rather than static inference, enabling adaptive refinement of hypotheses across stages of the drug discovery workflow. Complementing this, the platform adopts a tool-first architecture in which domain-validated computational methods—such as structure prediction, docking, ADMET modeling, and retrosynthesis planning—serve as the primary sources of scientific evidence.

These principles are operationalized through a supervisory orchestrator Agent, which translates high-level design objectives into coordinated multi-agent workflows. The orchestrator dynamically decomposes tasks, assigns them to specialized agents, and manages execution logic including dependency resolution, iteration, and convergence. Importantly, agent sequencing is not rigidly predefined but evolves based on intermediate results and emerging constraints, enabling context-aware adaptation of the workflow.

Together, ReAct-based execution and tool-first orchestration establish a unified framework in which reasoning, computation, and decision-making remain tightly coupled, supporting iterative refinement of molecular design strategies while maintaining clear linkage between intermediate evidence and downstream outcomes.

### Specialized Scientific Agents

To operationalize this framework, we implemented a suite of 26 specialized tools spanning structural biology, cheminformatics, ADMET prediction, and retrosynthesis. Each tool exposes structured inputs and outputs (e.g., SMILES strings, PDB IDs), enabling transparent orchestration across agents. Representative tools illustrating the multi-agent, tool-grounded architecture spanning structure analysis, molecular generation, property evaluation, synthesis planning, and workflow orchestration are presented in Table 1.

**Table 1:**
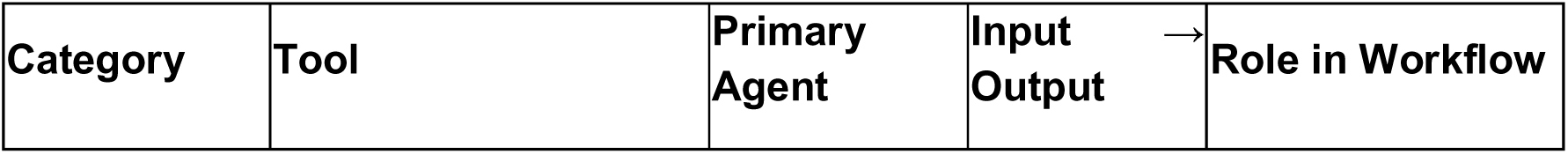
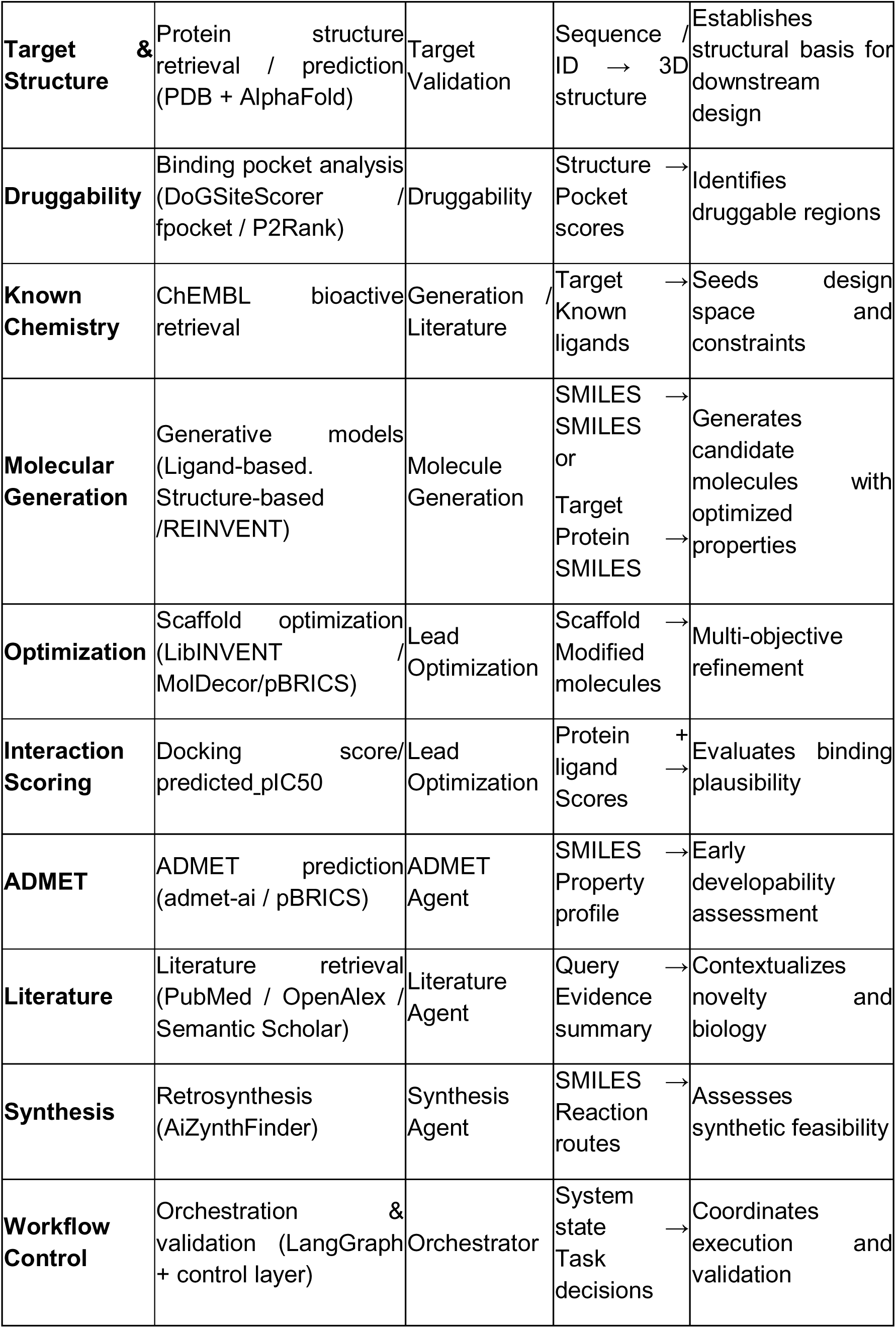
Representative tools illustrating the multi-agent, tool-grounded architecture spanning structure analysis, molecular generation, property evaluation, synthesis planning, and workflow orchestration.

### Target Validation Agent

Target validation is the first and foundational step of the multi-agent workflow, as all downstream design decisions critically depend on the availability, quality, and reliability of structural information for the biological target. The target validation agent can retrieve experimentally resolved protein structures from the Protein Data Bank (PDB)^19^ when available or generates high-confidence structural models using AlphaFold in cases where experimental structures are absent.^1^

The agent systematically assesses structural quality and uncertainty using confidence metrics such as predicted Local Distance Difference Test (pLDDT) scores, identifying poorly resolved regions that may impact reliability of binding-site identification. This early validation step prevents propagation of structural artefacts into later stages of molecular generation and prioritization, thereby reducing false confidence and downstream design risk.

### Druggability Assessment Agent

Following target validation, druggability assessment constitutes a critical early decision point that determines the presence of druggable binding pockets and whether it is feasible for the selected target. The druggability assessment agent systematically identifies and characterizes potential ligand-binding pockets on the validated protein structure and evaluates their suitability for small-molecule intervention. To ensure robust and consensus-driven assessment, the agent applies multiple complementary methods, including fpocket^20^, DogSiteScorer^21^, and P2Rank^22^, each of which captures distinct geometric, physicochemical, and evolutionary features associated with druggable binding sites. Based on this evidence, the agent determines whether the target should proceed to molecular design, whether an alternative design strategy is warranted, or whether the target should be deprioritized altogether.^23^ By elevating druggability assessment from a passive filtering step to an explicit decision-making process, this agent prevents premature commitment to structurally intractable targets and reduces downstream attrition in molecular generation and optimization stages.

### Molecule Generation Agent

Upon confirmation of target and pocket druggability, the molecule generation agent is responsible for proposing chemically viable candidate molecules through structure- and ligand-guided generative modeling.^3–8^ Molecular generation is performed using a combination of established open-source reinforcement learning frameworks and proprietary generative models previously developed by the authors. It can also incorporates the open-source tools like REINVENT framework for de novo and scaffold-constrained molecular design, alongside previously developed in-house generative models, namely ligand-based, structure-based, omic-based or synthesis-aware.^6–8,10^ Any other third-party tools can also be integrated through some easy steps. Molecule generation is formulated as an explicit multi-property optimization problem rather than a single-objective search. Reinforcement learning reward functions are constructed to balance multiple design criteria simultaneously, including predicted target-specific bioactivity, physicochemical properties, and other developability-related constraints including ADMET. This multi-objective optimization strategy follows previously described approaches for controllable molecular generation, enabling systematic exploration of trade-offs between competing objectives that commonly arise in early-stage drug discovery.^4^

Property evaluation and scoring within the reward formulation are driven by predictive models trained and validated in prior work.^4,8^ Importantly, the molecule generation agent does not function as an isolated design module. Generated molecules are continuously evaluated by downstream agents, including developability filtering, and synthetic feasibility analysis. Feedback from these agents is propagated back into subsequent generation cycles, allowing the reward function and optimization trajectory to be dynamically refined. This iterative, tool-grounded feedback enables progressive convergence toward chemically realistic and development-relevant candidates, while maintaining transparency over how specific molecular features are promoted or penalized during optimization.

### Lead Optimization Agent

Following initial molecule generation, the lead optimization agent is responsible for iterative refinement of candidate molecules to improve potency, selectivity, and overall developability while preserving core binding motifs. Rather than performing one-shot filtering, during lead optimization molecular structures are progressively modified based on quantitative feedback from structure- and property-based evaluations.

Chemical modifications are proposed using a combination of scaffold-aware generative models and targeted decoration strategies. In this particular work, the agent employs LibINVENT^24^ frameworks for scaffold-constrained optimization, alongside in-house developed MolDecor methodology^25^, which enables systematic exploration of substituent space around predefined molecular cores while maintaining medicinal chemistry relevance.^25^ These generative approaches allow focused optimization of lead series without disrupting established binding hypotheses. Similar to molecule generation, lead optimization is embedded within a multi-agent framework. Feedback from bioactivity, property prediction, and downstream developability assessments is propagated into subsequent optimization rounds, allowing the agent to adaptively refine design strategies.

### ADMET & Bioactivity

The prediction of ADMET properties helps to assess developability risks at an early stage, before significant resources are committed to predicted chemically or biologically optimal candidates. Recognizing that late-stage attrition in drug discovery is frequently driven by unfavorable pharmacokinetic or toxicity profiles, this agent systematically evaluates absorption, distribution, metabolism, excretion, and toxicity (ADMET) properties^13^ alongside predicted bioactivity in parallel with molecular design and optimization.

Importantly, ADMET assessment is embedded within the iterative agentic workflow rather than applied as a terminal filtering step. Predicted liabilities and risks are explicitly propagated upstream to inform subsequent molecule generation and lead optimization cycles, enabling targeted structural refinement rather than post hoc rejection. By shifting ADMET evaluation from a late-stage gatekeeping role to an early, decision-informing process, the agent reduces the likelihood of late-stage failures and supports the prioritization of chemically tractable, biologically relevant, and development-ready drug candidates. Therefore, ADMET & Bioactivity are included as tools in both the molecule generation and lead optimization agents.

### Literature agent

To ensure that molecular design decisions are grounded in prior biological and chemical knowledge, the literature & evidence agent systematically aggregates and contextualizes external evidence throughout the discovery workflow. As shown in **Fig. 1**, this agent operates in parallel with molecular generation and optimization, acting as an explicit evidence-integration layer rather than a post hoc validation step.

**Figure 1:**
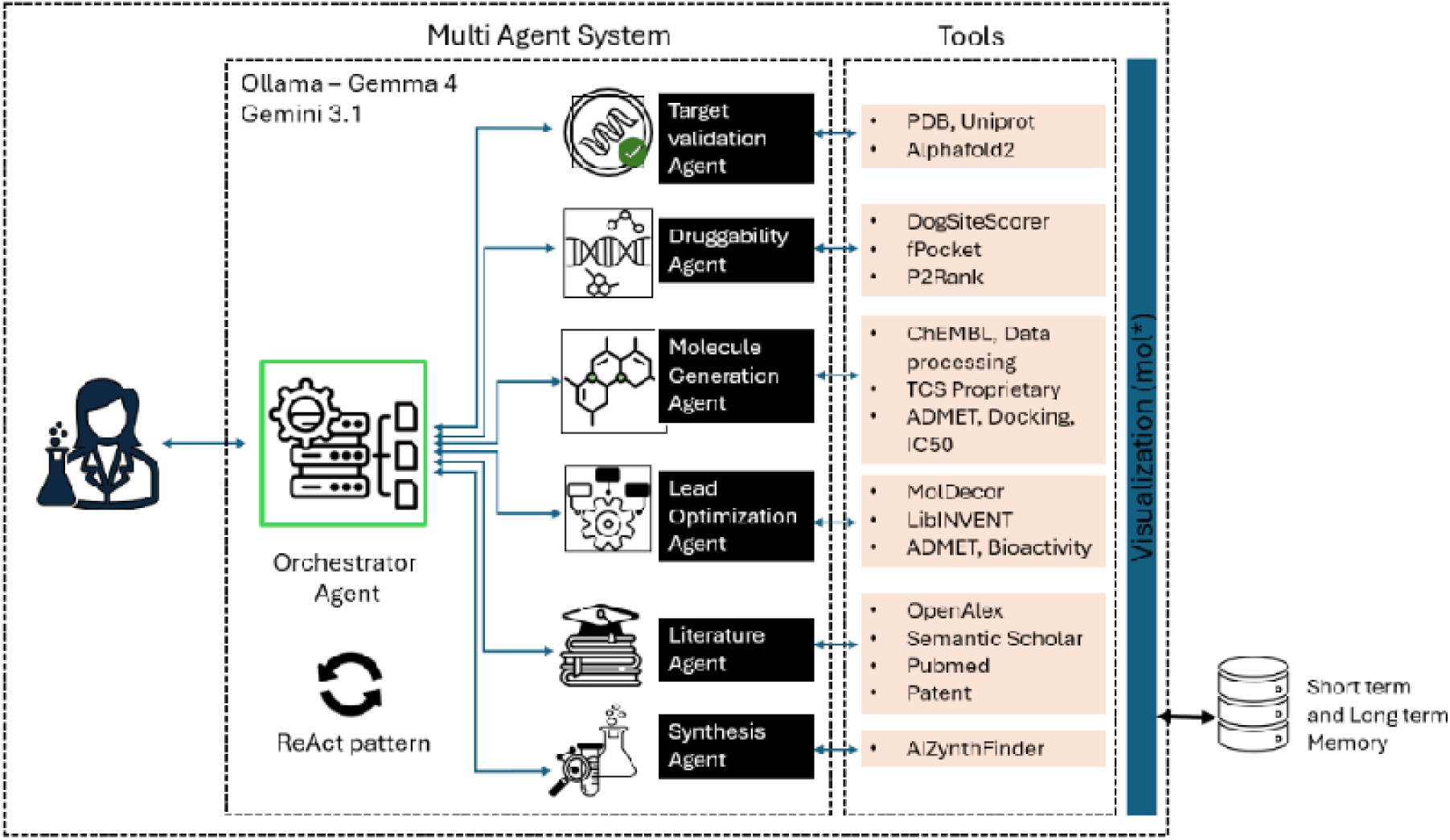
Overall architecture of TRACEDD, a tool-grounded multi-agent framework for reasoning-driven drug design. The workflow decomposes early-stage discovery into specialized scientific agents for target validation, druggability assessment, molecular generation, lead optimization, ADMET and bioactivity evaluation, literature evidence integration, and retrosynthesis. A supervisory orchestrator coordinates these agents through a Reason–Act–Observe loop, enabling explicit tool invocation, evidence-based decision-making, iterative feedback, and traceable rationale across the drug design cycle.

The agent mines relevant public and proprietary literature sources using OpenAlex^26^, Semantic Scholar^27^and PubMed to identify known ligands, reported bioactivity patterns, and previously observed liabilities associated with the target or related protein families. By explicitly mapping generated or optimized molecules to existing chemical and biological knowledge, the agent assesses novelty, previously reported limitations, and highlights areas where proposed designs overlap with or can be improved from known knowledge.

Rather than producing unstructured citations, the literature & evidence agent synthesizes this information into explainable evidence summaries that directly support downstream decisions. These summaries where literature-evidence supports or contradicts predicted activity or selectivity, and what uncertainties remain. Importantly, this evidence is propagated back to other agents through the shared system state (Fig. 1), allowing molecule generation, lead optimization, and prioritization steps to adapt based on earlier knowledge rather than implicit assumptions.

By embedding literature-driven reasoning directly into the agentic workflow, the system ensures that design decisions remain traceable, scientifically defensible, and aligned with existing knowledge, while still allowing systematic exploration beyond known chemotypes. This explicit integration of evidence reduces the risk of rediscovering well-established molecules, surfaces conflicts early, and supports transparent decision-making across iterative design cycles.

### Synthesis (Retrosynthesis) Agent

Synthetic feasibility is a critical yet frequently underestimated determinant of success in early-stage drug discovery. To address this, the synthesis (retrosynthesis) Agent explicitly evaluates synthetic tractability at an early stage, ensuring that proposed molecules are not only chemically and biologically promising but also practically synthesized. As illustrated in **Fig. 1**, this agent operates in parallel with molecular generation and lead optimization, contributing actionable constraints rather than acting as a late-stage rejection filter.

For each candidate molecule, the agent proposes viable synthetic routes using AI-driven retrosynthesis planning methods such as AiZynthFinder^28^, identifying plausible disconnections, reaction sequences, and intermediate accessibility. The feasibility of proposed routes is assessed based on factors such as reaction confidence, step count, availability of starting materials, and overall synthetic complexity. By including synthesis assessment in explicit route proposals, the agent provides transparent justification for why a molecule is considered tractable or impractical.

Importantly, the synthesis agent plays an active decision-making role within the agentic workflow. Molecules that exhibit attractive biological profiles but lack reasonable synthetic pathways are explicitly flagged or rejected, preventing chemically elegant but impractical designs from propagating downstream. Conversely, feedback from retrosynthetic analysis is propagated upstream to inform molecule generation and lead optimization, enabling structural modifications that improve synthetic accessibility without compromising core activity hypotheses.

In production settings, Gemini 3.1 Flash Lite is used as the default reasoning backbone because of its strong reasoning capability, low latency, and long-context support. Ollama-hosted Gemma 3 and Gemma 4 serve as privacy-preserving alternatives for deployments requiring complete on-premises execution.^29,30^ Together, these deployment options provide a flexible foundation for agentic drug-discovery workflows across a broad range of pharmaceutical R&D environments.

## Results

To evaluate the ability of TRACEDD to coordinate heterogeneous computational tools within a unified reasoning framework, we showed an end-to-end *de novo* drug design workflow targeting Janus kinase 2 (JAK2), a well-characterized kinase with extensive structural and bioactivity information. The objective was not simply to generate candidate molecules, but to assess whether the framework could progressively transform a high-level scientific objective into a series of evidence-linked decisions spanning target assessment, hypothesis generation, molecular design, optimization, prioritization, and synthesis planning.

### Target validation and protein structure retrieval/prediction

The discovery campaign began with target validation and structural assessment. Upon receiving JAK2 as the target, the target validation agent mapped the gene to its UniProt identifier and queried structural repositories (PDB) to identify available experimental structures. The framework retrieved 145 crystallographic structures and selected a high-resolution complex (PDB ID: 3UGC; 1.34 Å) as the starting point for downstream analysis. This process was fully traceable through the reasoning log, which explicitly recorded target mapping, structure retrieval, and structure selection criteria (Fig. 2).

**Figure 2:**
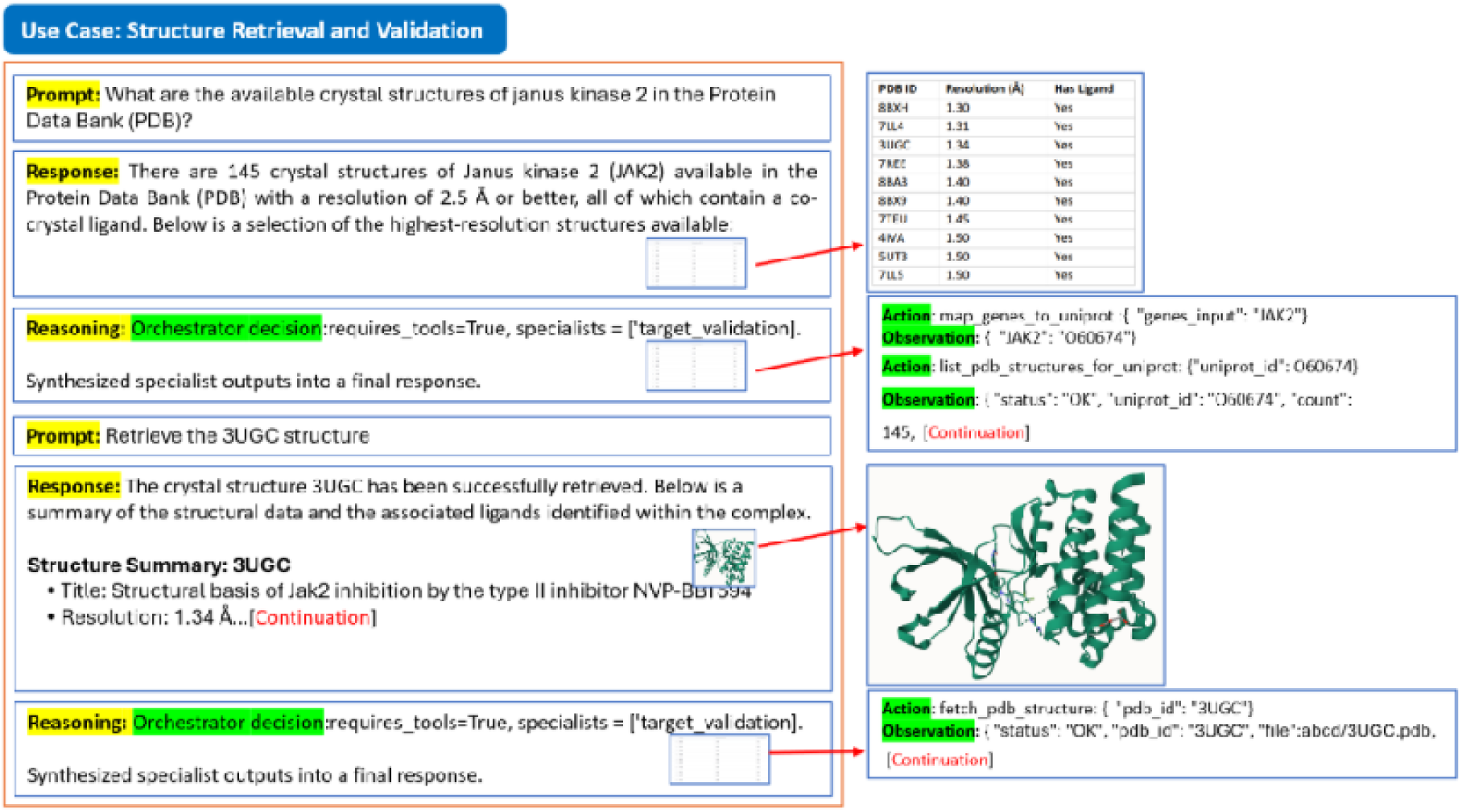
Adaptive target validation workflow for protein structure acquisition in TRACEDD. The Target Validation Agent maps the user-specified target to a unique protein identifier, queries experimental structural repositories such as the PDB, and retrieves available high-resolution structures when present. If no suitable experimental structure is available, the workflow transitions to AlphaFold-based structure prediction, ensuring that downstream structure-based design proceeds using the best available structural representation while preserving data provenance and reasoning traceability.

To demonstrate adaptability under differing data availability conditions, the framework was additionally evaluated to check if it can work on targets lacking experimental structure. In this scenario, the agent automatically invoked AlphaFold-based structure prediction and incorporated the resulting model into the same downstream workflow used for experimentally resolved structures (Fig. 3). This conditional behavior illustrates how TRACEDD dynamically adapts its workflow while preserving continuity of analysis.

**Figure 3:**
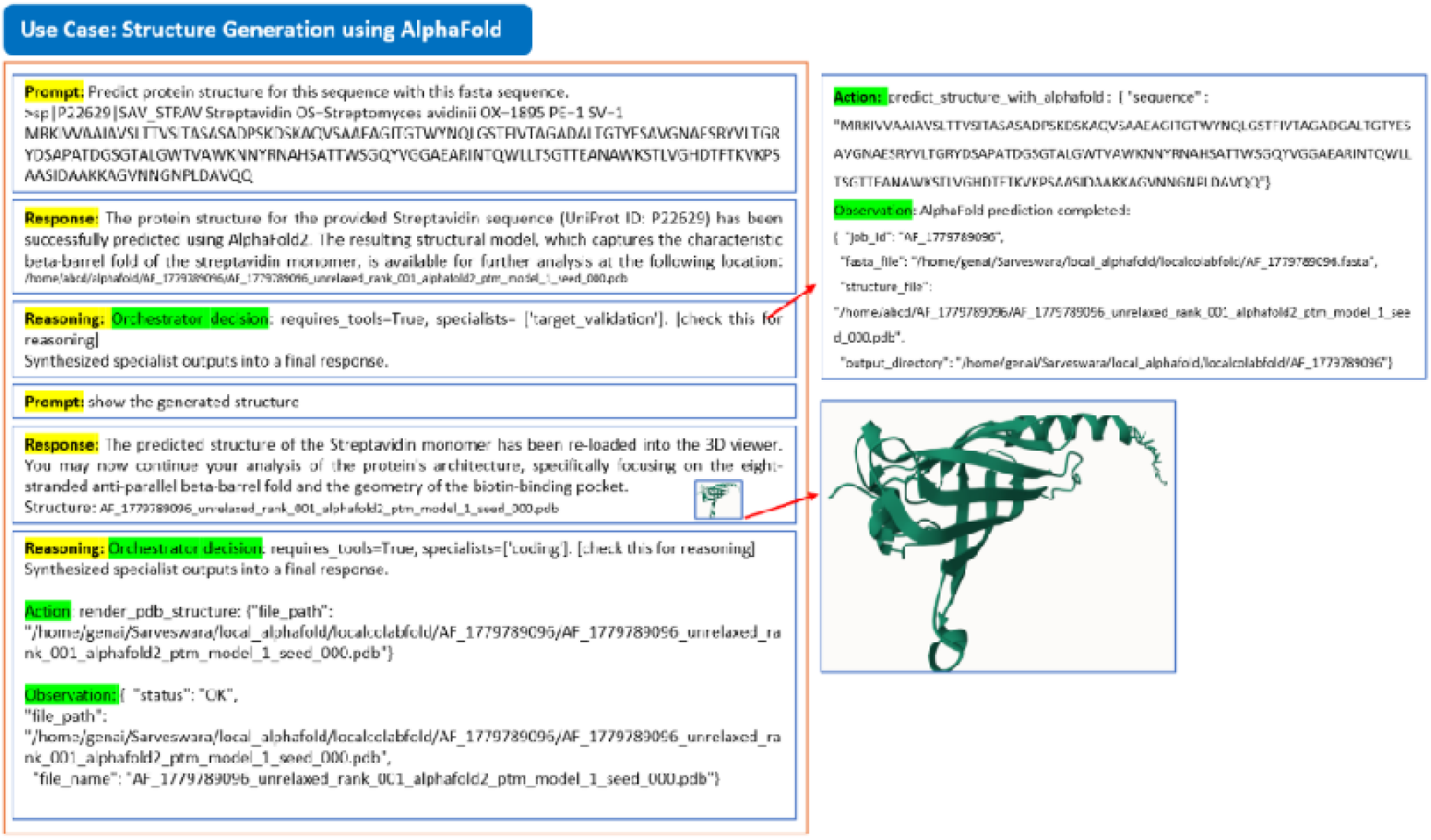
Representative structure retrieval and prediction outcomes generated by the Target Validation Agent. For targets lacking experimental structures, the agent invokes AlphaFold-based modeling and integrates the predicted PDB structure into the same downstream workflow, enabling consistent visualization, pocket analysis, and structure-based drug design.

Conversion of holo complexes into ligand-free receptor structures is often required for next subsequent steps. Consequently, the framework delegated apo-structure generation to a specialist coding agent. The agent programmatically removed ligand-associated entries from the JAK2 crystal structure and generated a cleaned apo receptor representation suitable for pocket analysis, docking, and molecular design (Fig. 5). This example highlights how TRACEDD bridges high-level scientific intent and executable computational operations through the same agentic reasoning framework.

### Agent-Guided Hypothesis Generation Prior to Molecular Design

Rather than initiating molecular generation immediately, here TRACEDD was first instructed to construct an explicit design hypothesis intended to constrain and guide subsequent exploration of chemical space. To achieve this, the orchestrator coordinated to understand target validation, literature retrieval, bioactivity analysis, and property-analysis (Fig. 4).

**Figure 4:**
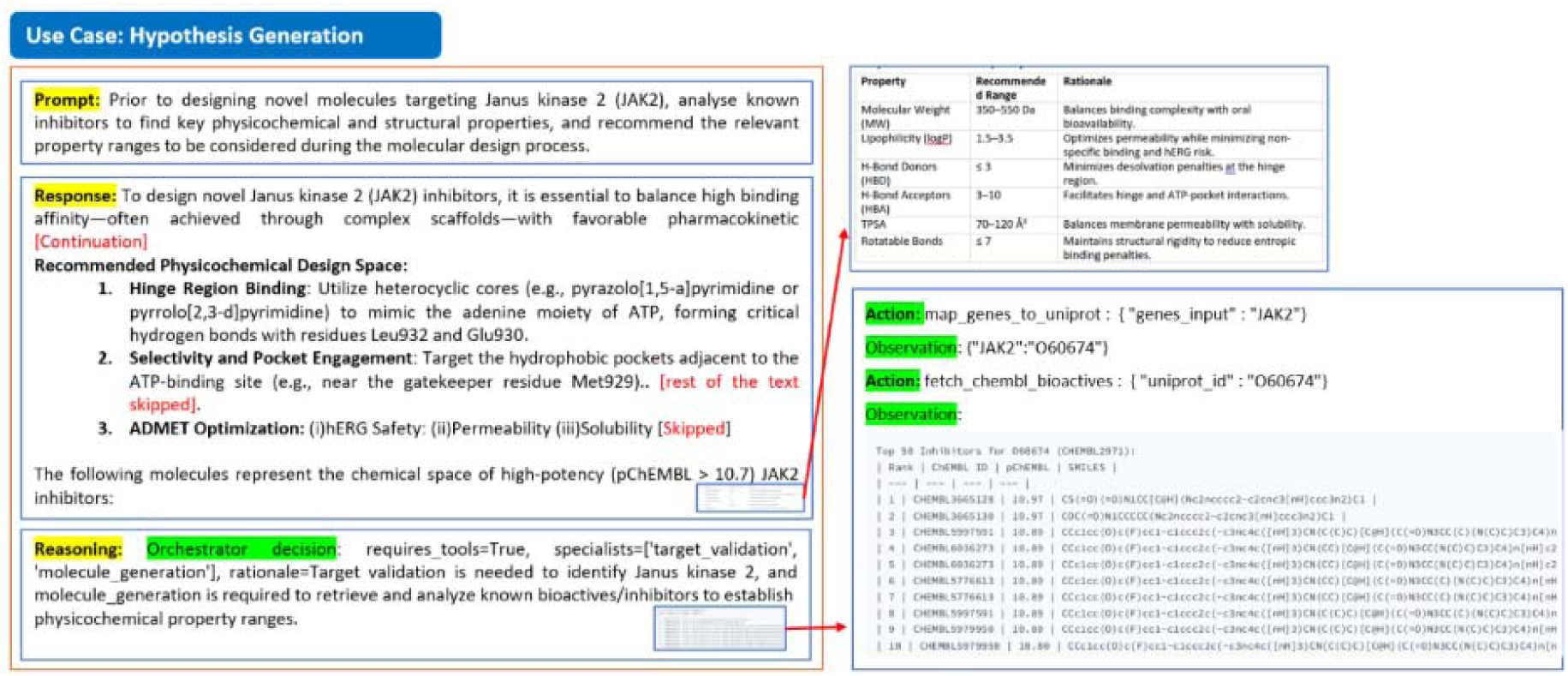
Agent-guided hypothesis generation for JAK2 inhibitor design prior to molecular generation. The orchestrator coordinates target validation, retrieval of potent known JAK2 bioactives, physicochemical and ADMET profiling, structural motif analysis, and evidence synthesis to define a mechanistically grounded design hypothesis. The resulting design space captures preferred kinase-binding scaffold features, hinge-region interaction motifs, selectivity considerations, property ranges, and developability constraints for downstream closed-loop molecule generation.

Known potent JAK2 inhibitors were retrieved from curated bioactivity data sources (e.g. ChEMBL), focusing on highly potent inhibitors with pChEMBL cutoff. Analysis of these potent inhibitors showed that high JAK2 activity is commonly associated with moderately lipophilic, scaffolds that occupy the ATP-binding cleft and extend into adjacent hydrophobic regions. Across representative compounds, the agent identified a relatively consistent physicochemical profile compatible with oral drug-like behavior. Although several high-affinity molecules approached the upper range of molecular complexity, the derived design space emphasized the need to balance binding strength with solubility and permeability, rather than optimizing potency alone. These observations were converted into explicit property ranges and design constraints to support downstream molecular generation.

The agent further analyzed structural motifs associated with existing JAK2 inhibitors by comparing ligand scaffolds with conserved kinase binding modes. A recurring feature was the presence of heterocyclic cores capable of mimicking the adenine moiety of ATP and forming canonical hinge-region hydrogen bonds with residues such as Leu932 and Glu930. Scaffold classes such as pyrazolo[1,5-a]pyrimidines and pyrrolo[2,3-d]pyrimidines emerged as commonly observed scaffolds within the potent inhibitor set. Beyond hinge binding, the agent highlighted engagement of hydrophobic pockets near the gatekeeper residue Met929 as an important strategy for improving JAK2 selectivity. Because the ATP-binding site is highly conserved across kinases, the system emphasized the importance of exploiting subtle structural features, such as P-loop orientation and DFG-in conformations, rather than relying only on bulky substituents for selectivity.

In parallel, the framework evaluated developability-related issues from representative high-potency inhibitors. Predicted ADMET properties indicated that most molecules satisfied baseline drug-likeness criteria, but several compounds showed lipophilicity-associated liabilities, particularly potential hERG risk. In the current use case, default properties such as lipophilicity, hERG liability, permeability, and solubility were considered, although the framework can be adapted to include additional endpoints depending on user requirements. Importantly, these ADMET properties were not treated as late-stage filters (Fig. 4). Instead, the agent incorporated them directly into the design hypothesis by recommending strategies such as limiting excessive hydrophobic surface area and positioning polar functionalities toward solvent-exposed regions.

Importantly, these observations were not retained as descriptive summaries. Instead, TRACEDD converted them into explicit design constraints that were propagated to downstream molecular generation agents. This transformed heterogeneous structural, chemical, and pharmacological evidence into an actionable and mechanistically grounded search space (Fig. 4). The resulting workflow demonstrates how agentic reasoning can formalize scientific hypothesis generation as an intermediate step between data collection and molecular design.

### Code agent-mediated generation of ligand-free (apo) protein structures

To evaluate whether the agentic framework can execute precise, programmatic structural transformations required for structure-based drug discovery, we next examined a use case involving automated generation of a ligand-free (apo) protein structure from an experimentally resolved holo complex (Fig. 5).

**Figure 5:**
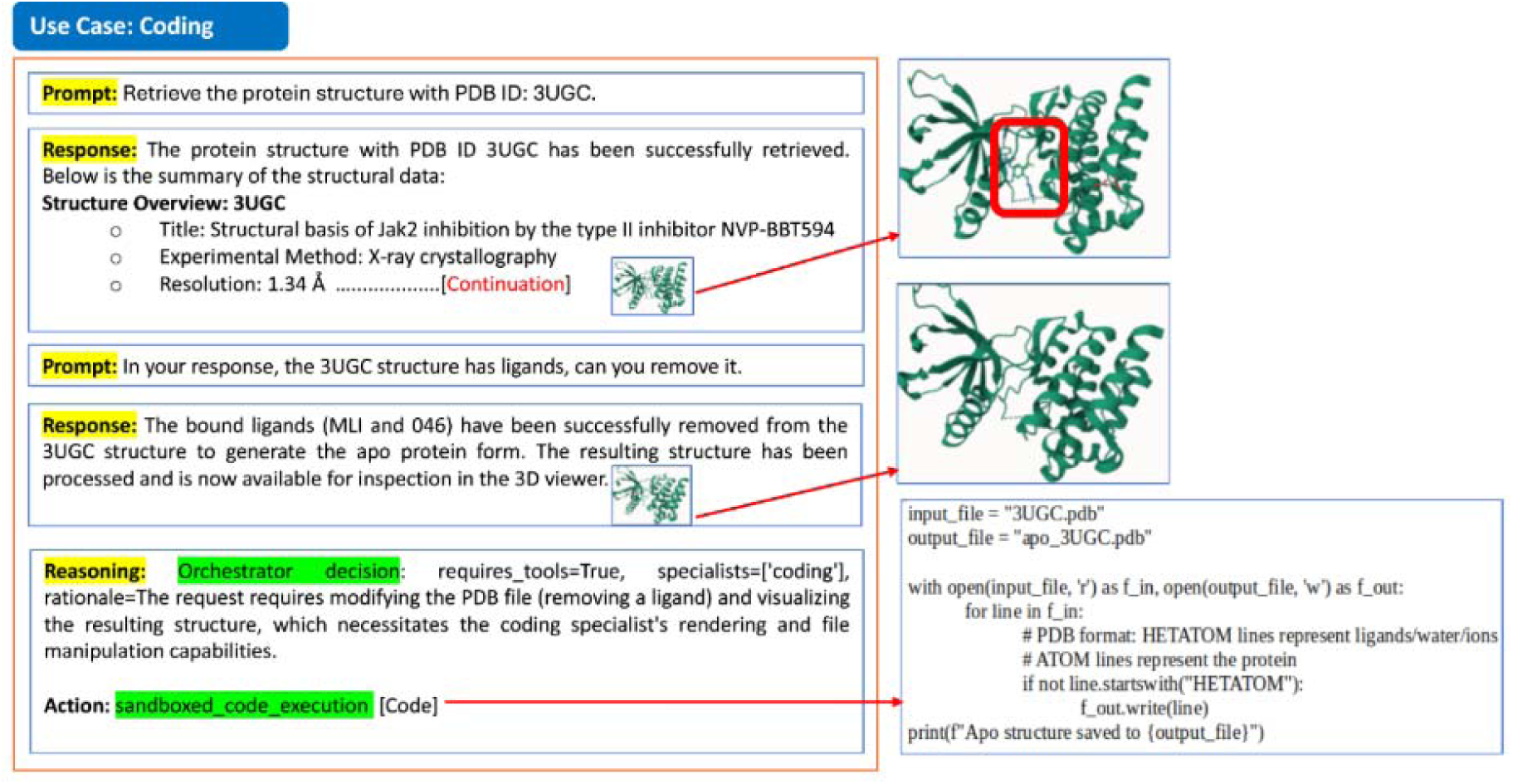
Code-agent-mediated generation of a ligand-free apo JAK2 structure from a holo crystal complex. Starting from the experimentally resolved JAK2 structure 3UGC, the framework identifies co-crystallized ligand and non-protein records, delegates the transformation to a coding specialist agent, and programmatically removes ligand-associated entries while retaining the protein structure. The resulting apo_3UGC.pdb file provides a reproducible, ligand-free structural artifact suitable for pocket analysis, docking, and downstream structure-based drug design.

Upon specification of a target structure (PDB ID: 3UGC), the agent retrieved the corresponding high-resolution X-ray crystal structure (1.34 Å), which captures JAK2 bound to a inhibitor. Initial parsing of the structural metadata identified the presence of co-crystallized small-molecule ligands within the file, necessitating their removal prior to downstream applications such as pocket analysis, docking, or conformational sampling.

The task of converting the holo structure into an apo form was delegated by the orchestrator to a coding specialist agent, reflecting an explicit decision that this request required executable manipulation rather than static analysis. The coding agent programmatically processed the structure file by selectively retaining protein ATOM records while removing ligand and non-protein HETATM entries. This transformation resulted in a cleaned apo protein structure, preserved in standard PDB format.

The resulting ligand-free structure (apo_3UGC.pdb) was rendered directly within the workspace, enabling immediate visual inspection and validation (Fig. 5). Importantly, this process did not rely on pre-stored apo templates or manual intervention, but instead executed a reproducible, code-driven structural edit derived from the user’s intent. The reasoning trace captured by the system explicitly documents both the need for tool execution and the resolution of file-level dependencies, illustrating the agent’s ability to recover from intermediate execution errors and adapt to the underlying file system state.

This use case demonstrates that the agentic system can bridge high-level scientific intent (“generate an apo structure”) with low-level computational operations, producing artifacts that are immediately usable in subsequent structure-based workflows. By embedding executable coding actions within the same agentic loop used for scientific reasoning, the framework enables seamless transition between conceptual reasoning and concrete molecular data transformation.

### Agent-driven drug design with RL

Using the validated receptor structure and agent-derived design hypothesis, TRACEDD subsequently executed a *de novo* design workflow. The Druggability Agent first characterized ligand-binding pockets and identified the ATP-binding region as the most promising site, assigning a high druggability score and confirming its suitability for small-molecule intervention (Fig. S1).

The molecule generation agent then initiated a *de novo* design using reinforcement learning (RL)-based molecular generation. Unlike conventional virtual screening approaches, which are constrained by predefined libraries, the framework explored novel chemical space conditioned on both the structure the target pocket (based on docking) and the design constraints established during the hypothesis-generation phase. During RL optimization, the pIC50 score (calculated based on top docking pose) was used as one of the rewards, such that molecules predicted to bind more favorably to the target pocket received higher rewards. This formulation enables the model to iteratively adjust molecular structures toward improved predicted binding affinity where the chosen reward (binding affinity, solubility etc.) drives molecular optimization toward pocket-specific binding hypotheses, providing a systematic pathway from protein structures to prioritized candidate molecules.

**Figure 6:**
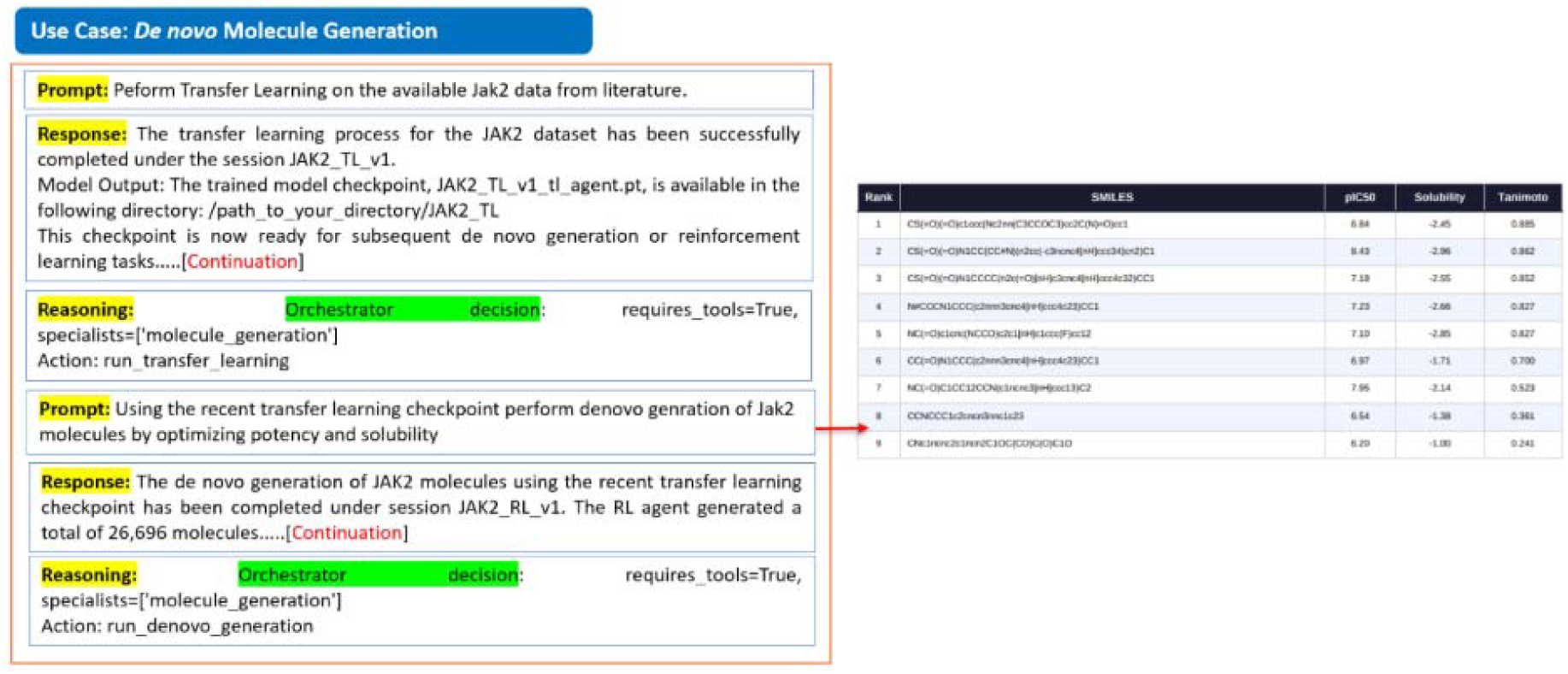
*De novo* molecular generation and multi-property optimization implemented using REINVENT4. Transfer learning (TL) adapts the generative model toward the chemical space of known JAK2 inhibitors, followed by reinforcement learning (RL) to optimize the generated molecules against the predefined activity, physicochemical, and developability objectives. The workflow yields novel candidate molecules with improved target-relevant property profiles.

### Agent-Guided Candidate Selection and Synthesis Planning

To prioritize candidates from the *de novo* generated set, the framework first analyzed known JAK2 inhibitors to identify recurring functional groups and molecular fragments associated with established target activity (Fig. 7). These target-relevant motifs were then encoded as substructure queries and systematically searched across the generated molecules. For each candidate, the agent recorded the presence of individual motifs and calculated the total number of matched fragments, providing an interpretable link between the newly generated structures and the known JAK2 inhibitor chemical space.

**Figure 7:**
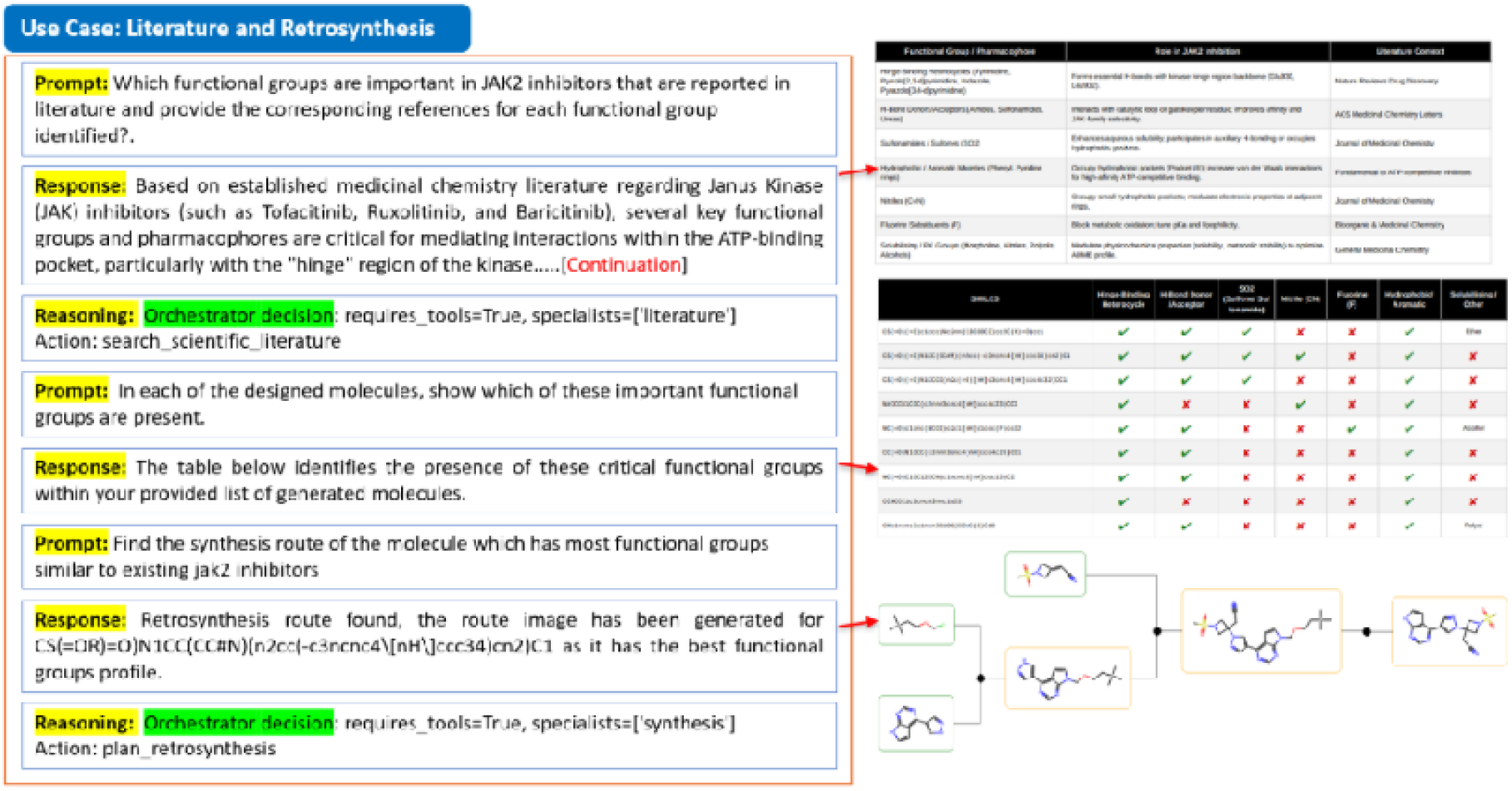
Functional-group-based prioritization of de novo generated JAK2 inhibitor candidates. Relevant functional groups and fragments are first identified from existing JAK2 inhibitors and then searched within the generated molecules. The table summarizes fragment matches across candidates, enabling prioritization of the molecule containing the maximum number of JAK2-relevant fragments. For the prioritized molecule, the Synthesis Agent subsequently uses AiZynthFinder to propose a plausible retrosynthetic route and assess synthetic feasibility.

The fragment-match shown in **Fig. 7** enables transparent comparison of the generated candidates and supports prioritization of the molecule containing the maximum number of JAK2-relevant fragments. This criterion complements the activity, physicochemical, and developability objectives used during REINVENT4 generation by retaining candidates that combine optimized predicted properties with recognizable target-associated chemical features.

The prioritized molecule was subsequently passed to the synthesis agent for retrosynthetic assessment using AiZynthFinder. The agent used the candidate structure as input to search for plausible disconnections and multistep routes leading to accessible precursors, thereby providing an explicit synthesis hypothesis alongside the molecular prioritization result. Integrating fragment-level evidence with retrosynthetic planning allows the framework to progress from de novo generation to a candidate that is both supported by known JAK2 chemical features and accompanied by a computationally proposed route for experimental evaluation.

## Discussion and Conclusion

The TRACEDD framework demonstrates that agentic AI can support end-to-end drug discovery workflow execution by coordinating specialized computational tools within a unified reasoning framework. Across the JAK2 case studies, the system successfully integrated target validation, hypothesis generation, structure preparation, molecular design, lead optimization, and synthesis planning into a continuous evidence-driven workflow. Rather than focusing solely on molecule generation, the results illustrate how agentic systems can transform high-level scientific objectives into a sequence of traceable and computationally grounded decisions.

A central contribution of TRACEDD is its treatment of reasoning as an explicit component of the discovery process. Unlike conventional computational pipelines that expose only inputs and outputs, the framework preserves reasoning traces linking observations, hypotheses, tool invocations, and downstream decisions. Consequently, decisions such as pocket selection, design constraint identification, candidate prioritization, and synthesis planning remain connected to verifiable computational evidence. Furthermore, workflow execution adapts dynamically to intermediate results, enabling the system to switch seamlessly between alternative strategies such as experimental structure retrieval, structure prediction, or code-based molecular manipulation. This adaptive behavior more closely reflects how scientific discovery proceeds in practice, where hypotheses are continuously refined in response to emerging evidence.

The results also reinforce the value of a tool-first paradigm for scientific AI. Within TRACEDD, large language models function as reasoning and orchestration layers, whereas domain-specific tools remain responsible for quantitative prediction and validation. This separation is important because the strengths and limitations of LLMs differ fundamentally from those of scientific models. LLMs excel at integrating heterogeneous information, generating hypotheses, and coordinating workflows, whereas tasks requiring quantitative accuracy, mechanistic consistency, or adherence to physical constraints are more appropriately handled by specialized computational methods. By grounding decisions in validated tools, the framework reduces dependence on latent model inference and maintains a transparent link between conclusions and supporting evidence.

An important observation emerging from this work is the role of hypothesis generation as an intermediate layer between data acquisition and molecular design. Rather than treating molecular generation as an unconstrained optimization problem, TRACEDD first constructs explicit design hypotheses from structural, chemical, biological, and developability evidence. These hypotheses define a mechanistically informed search space that guides downstream optimization and candidate selection. This finding suggests that the greatest value of agentic AI in drug discovery may lie not in autonomous molecule generation itself, but in its ability to generate, refine, and test scientific hypotheses across the discovery lifecycle.

There are several limitations as well. The current study focuses on representative use cases and therefore emphasizes functional capability rather than large-scale benchmarking. In our future work, we aim to establish quantitative evaluation frameworks for agentic scientific systems, including measures of task completion, reasoning consistency, tool-selection accuracy, recovery from execution failures, and agreement with experimental observations. In addition, human expertise remains essential for interpreting biological relevance, assessing structural uncertainty, and selecting candidates for experimental validation.

Overall, TRACEDD supports a broader shift from workflow automation toward evidence-linked scientific reasoning. By combining agentic orchestration with tool-first computational analysis, the framework demonstrates how reasoning, computation, and decision-making can remain distinct yet tightly connected throughout drug discovery. Such an approach offers a pathway toward more transparent, and interpretable AI-assisted molecular design.

## Supplementary Material

### Memory & Knowledge Layer

The agentic framework incorporates a unified memory and knowledge layer that maintains continuity, traceability, and cumulative learning across drug discovery workflows. Complementary short-term and long-term memory components support iterative reasoning within a session and preserve knowledge across programs. Short-term memory is maintained within the execution state (e.g., LangGraph), capturing the full sequence of agent interactions, tool invocations, intermediate outputs, and reasoning traces during a single workflow. This ensures that agents can iteratively refine hypotheses, revisit prior results, and propagate uncertainty across stages without loss of context. Each decision step is explicitly linked to the corresponding tool call and observed output, preserving a structured trace of the reasoning process.

Long term memory persisted using a lightweight SQLite database, which serves as the system’s knowledge store. In addition to checkpointing validated scientific outputs (e.g., structural templates, prioritized scaffolds, ADMET profiles, retrosynthetic routes), the SQLite layer also stores user sessions and past chat conversations. This design ensures that both scientific evidence and conversational context can be recalled in future sessions, enabling continuity of dialogue, reproducibility of reasoning, and cumulative knowledge across projects. By integrating these two layers, the system captures both transient reasoning dynamics and accumulated scientific knowledge. Historical design trajectories, failure modes, and successful optimization strategies can be retrieved and reused in subsequent campaigns, reducing redundant exploration and improving consistency in decision-making. This design mirrors human discovery processes, where iterative discussions inform immediate actions while institutional knowledge guides long-term strategy. Overall, the Memory and Knowledge Layer ensures that agentic workflows remain context-aware during execution and knowledge-rich over time, supporting adaptive, evidence-driven drug design with persistent traceability of decisions and outcomes.

### Technical Architecture: LLM Backends and Orchestration

The platform’s technical foundation combines cloud-based inference, local deployment options, and a multi-agent orchestration framework to balance reasoning quality, cost efficiency, and data governance requirements.

### Google Gemini API (Primary Backbone LLM)

The primary reasoning backbone is Google’s Gemini 3.1 Flash Lite model, accessed via the Google AI API and integrated through the langchain_google_genai package. Gemini 3.1 Flash Lite is a multimodal transformer-based large language model with native tool-calling support and an extended context window of up to 1 million tokens, enabling reasoning across long scientific workflows. Within the application, the model operates deterministically (temperature = 0) and is bound to the full set of 26 LangChain Structured Tool objects. Function calls are executed via LangGraph, which appends tool outputs back into the conversation state for iterative reasoning. Authentication is handled via environment variables, and pricing remains economical (∼$0.10 per million input tokens, ∼$0.40 per million output tokens). A typical drug discovery session costs under $0.10, making Gemini Flash Lite highly competitive relative to GPT-4-class models.

### Ollama + Gemma 3 and Gemma 4 (Local LLM Alternatives)

For users requiring full data privacy or cost-free inference, the system supports local deployment via Ollama. Ollama is an open-source inference server that hosts quantized models in GGUF format and exposes an OpenAI-compatible REST API. Two local backbones are supported:

- Gemma 3 27B Instruct: A 27-billion-parameter instruction-tuned model developed by Google DeepMind, optimized for structured reasoning and synthesis. Running in half-precision (FP16), it requires ∼36–40 GB of GPU VRAM, achievable on NVIDIA A100 or A6000 GPUs.
- Gemma 4 Instruct: The latest generation, offering improved reasoning quality and efficiency compared to Gemma 3 ^30^. Deployed via Ollama, Gemma 4 ^31^ provides a fully offline, zero-API-cost alternative while ensuring sensitive molecular structures and proprietary datasets remain entirely within local environment.

These local deployments eliminate per-token API costs and guarantee that no proprietary data leaves infrastructure. The trade-offs include higher latency (2–10 seconds per generation vs. ∼0.5 seconds for Gemini Flash Lite), hardware requirements, and rare quality gaps in specialized cheminformatics reasoning compared to Gemini’s larger-scale training. However, for regulated pharmaceutical settings, the privacy and cost benefits are substantial.

### LangGraph Agent Orchestration Framework

Multi-agent orchestration is implemented in LangGraph, a directed-graph framework for stateful LLM applications. Each node represents a specialist agent (e.g., target validation, ADMET profiling, synthesis planning), while edges define routing logic. The agent state is maintained as a Python TypedDict containing the full message history. LangGraph supports streaming outputs, checkpointing, and conditional routing, ensuring workflows remain auditable and reproducible. Every tool invocation, input, and output is logged, enabling transparent reasoning and human-in-the-loop validation.

### Comparative deployment architectures

TRACEDD supports both cloud-hosted and on-premises deployment configurations, enabling adaptation to diverse pharmaceutical research and development environments. The primary cloud deployment utilizes Gemini 3.1 Flash Lite, whereas local deployments are supported through Ollama-hosted Gemma 3 and Gemma 4 models. These deployment modes present distinct trade-offs in reasoning capability, latency, operational cost, governance, and infrastructure requirements.

Cloud-based deployment offers rapid inference, minimal operational overhead, and access to large-context reasoning capabilities. In our implementation, Gemini 3.1 Flash Lite provides low-latency responses and supports long-context scientific workflows, making it well suited for interactive discovery sessions and complex multi-step reasoning tasks. Because model hosting, maintenance, and scaling are managed externally, users can access advanced reasoning capabilities without the need for dedicated model infrastructure.

In contrast, local deployment through Ollama prioritizes data sovereignty, regulatory compliance, and operational control. By executing all inference within the customer environment, sensitive molecular structures, proprietary biological data, and project-specific knowledge remain entirely on-premises. This approach eliminates recurring API costs after infrastructure provisioning and enables integration within highly regulated pharmaceutical environments where external data transmission may be restricted. Although local models generally exhibit higher inference latency and may not consistently match the reasoning performance of larger cloud-hosted models, recent generations such as Gemma 4 substantially narrow this gap while preserving the advantages of privacy-preserving deployment.

The choice between deployment modes therefore depends on organizational priorities. Cloud-hosted deployments favor reasoning performance, scalability, and ease of operation, whereas local deployments emphasize data governance, auditability, and cost predictability. By separating orchestration from model backends, TRACEDD allows the same agentic workflow to operate across both environments without modification to the underlying scientific logic. This flexibility enables organizations to balance performance, privacy, compliance, and operational considerations according to the requirements of specific discovery programs.

**Figure S1:**
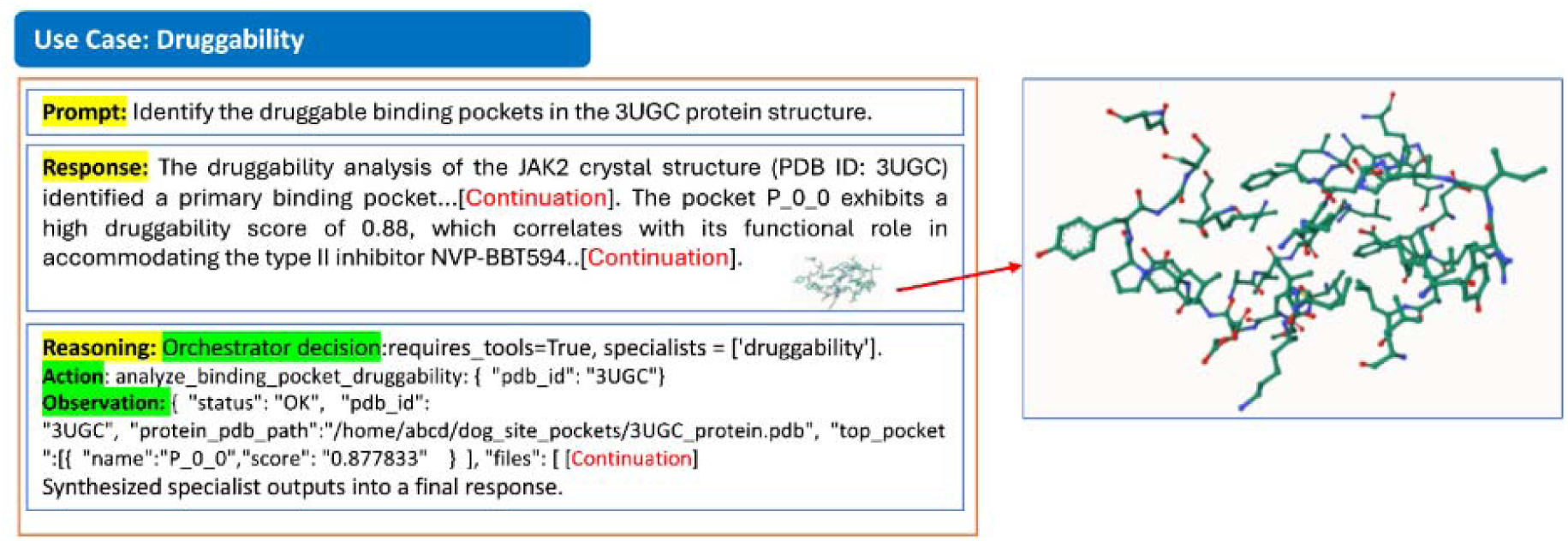
Druggability Agent-mediated identification and ranking of putative binding pockets in the JAK2 receptor structure using the DoGSiteScorer package. The agent evaluates predicted pockets using geometric and physicochemical descriptors and ranks them according to their druggability scores, supporting selection of the highest-ranked pocket for downstream docking and structure-based molecular design.

